# Cell-free pathway prototyping enables cost-effective biomanufacturing of 1,2,4-butanetriol at the 1-L scale

**DOI:** 10.64898/2026.08.03.742631

**Authors:** Blake J. Rasor, Katherine A. Rhea, Isaiah D. Richardson, J. Tyler Lazar, Marilyn S. Lee, Ethan N. Walters, John R. Biondo, Frank J. Kragl, David C. Garcia, Kyle Zolkin, John P. Davies, Matthew W. Lux, Ashty S. Karim, Michael C. Jewett

**Author notes:** These authors contributed equally to this work (Blake J. Rasor, Katherine A. Rhea). **Correspondence** Matthew W. Lux, Ashty S. Karim, Michael C. Jewett.

## Abstract

Biomanufacturing offers sustainable alternatives to chemical synthesis under lower temperatures and pressures than traditional catalytic methods. However, the slow pace and iterative engineering bottlenecks of cell strain development restrict the feasible biological design space. Cell-free systems circumvent these constraints, providing a flexible and high-throughput screening approach to accelerate pathway prototyping and enzyme optimization but are not typically used for manufacturing scale-up. To understand the scalability of cell-free biosynthesis, we establish an end-to-end fully cell-free architecture to discover, develop, and scale the biosynthesis of 1,2,4-butanetriol (BT), a high-value industrial platform chemical. First, we systematically screened ~150 enzymes across the 4-step pathway from xylose to BT to identify highly active homologs for each reaction. Next, we applied statistical Design of Experiments to optimize reaction formulations for cost and titer. Finally, the maximum-titer and minimum-cost formulations were scaled up across five orders of magnitude, from 10-µL to 1-L reactions. This resulted in peak volumetric productivities of ~1 g/L/h and yields over 13 g of BT in a single 1-L reaction, with raw substrate costs totaling just $3.00 per liter. This work expands the diversity of enzymes tested for BT synthesis and establishes a blueprint for advancing industrial biochemical manufacturing fully *in vitro*.

## Introduction

Biomanufacturing offers a sustainable route for chemical synthesis, with significant investments from both the public and private sectors recognizing the burgeoning potential of biotechnology^1–3^. Biological processes to manufacture molecules ranging from flavors and fragrances to fuels and pharmaceuticals have been developed both in cells and in cell-free systems comprising purified enzymes or crude cell extracts. Key examples include acids^4,5^, terpenes^6,7^, alkaloids^8^, and other natural products^9,10^. The most valuable compounds for biomanufacturing can be chemically converted to multiple downstream products with diverse applications^11,12^, enabling access to economies of scale. One such platform chemical is 1,2,4-butanetriol (BT), which serves as a highly functional building block for fuels and materials with potential for chemical modifications at the three hydroxyl groups^13–15^.

Conventional methods for BT synthesis modify fossil-derived malate via (i) borohydride reduction which generates stoichiometric excess of borate waste products^16^ or (ii) catalytic hydrogenation which requires elevated temperatures and high pressures of hydrogen^17^. The inherent drawbacks of these chemical synthesis routes have restricted market expansion, so metabolic engineers have explored pathways for biomanufacturing BT using milder, waste-minimized reaction conditions^13^. The 4-carbon triol is biologically accessible at high yields (>40 g/L) from multiple metabolic routes in a range of microorganisms and purified enzymes^14,15^. While metabolic routes from both 5- or 6-carbon sugars have been validated, the conversion of xylose to BT has been extensively explored due to high yields and the ability to use sustainable, xylose-rich substrates such as lignin hydrolysates^12,15,18^.

The highest yielding BT biosynthesis process reported to date used cell-free systems^14^, which offer advantages including enhanced physiochemical control, higher volumetric productivities, and increased toxicity thresholds compared to cellular fermentations^9,19–24^. However, the scalability of cell-free reactions has been historically limited by the product-to-substrate cost ratios^25–27^, due in part to the absence of self-regeneration mechanisms for enzymes and cofactors. Despite this limitation, cell-free gene expression (CFE) can successfully scale for high-value protein production in both bacterial and plant-based extracts^28,29^. Bulk chemicals, however, require significantly cheaper bioprocesses for economic viability among petrochemical competitors^4,5,25,30^. This reality led to the use of cell-free reactions for microliter-scale prototyping efforts to identify the best sets of enzymes for biochemical synthesis^31–33^. Key innovations from these efforts include successful identification of high-yielding biosynthetic pathways for butanol^34^, muconic acid^35^, hexanoic acid^36^, and terpene synthesis^7^, accelerated strain engineering by screening gene knockout candidates^37^, and the development of a C1 conversion pathway in the context of *E. coli* lysate^38^. Insights from these prototyping efforts along with demonstrations of cell-free biomanufacturing to synthesize high titers of terpenoids or isobutanol within bench-scale bioreactors up to 15 mL establishes a growing potential for economically scaling cell-free biochemical synthesis^9,19^.

In this work, we report an end-to-end pipeline for discovery, development, and manufacturing of biochemicals in cell-free systems, demonstrated with 1,2,4-butanetriol biomanufacturing. First, we use small-scale reactions to rapidly discover preferred enzymes for the 4-step pathway from xylose to BT from 141 enzyme variants. Next, we develop optimized reaction composition using machine learning-enabled design-of-experiments to minimize the cost and complexity while maximizing BT titers. A minimal formulation with reagent costs less than $3 per liter (excluding cell lysate) remained capable of producing 9 g/L BT during validation at small scale. Finally, we scaled cell-free BT synthesis 5 orders of magnitude from 10 µL to 1 L, achieving titers over 10 g/L with optimized formulations. This streamlined, purification-free manufacturing paradigm establishes a framework for cell-free biomanufacturing beyond established laboratory prototyping workflows.

## Results

### Discovering highly active enzyme variants for BT synthesis

Screening for enzymes with robust kinetics and high substrate affinity can improve biosynthetic pathway performance and alleviate bottlenecks. In previous synthesis of BT, relatively few enzymes variants were explored in a range of cells and purified cell-free systems^13^. Yet, the potential for further pathway improvement by identifying high-performance enzymes through compressed design-build-test-learn (DBTL) cycles offered by cell-free gene expression (CFE) warrants further screening efforts^39–43^. Here, we used CFE to establish a reference set of enzymes, validate cell-free BT synthesis, and then identify an optimized set of enzymes.

BT biosynthesis from xylose requires four enzymatic steps (**Fig. 1A**). In an initial enzyme search, we identified 2-3 enzyme families for each of the four steps in the pathway, including NAD- and NADP-dependent variants for steps 1 and 4 (**Fig. 1A**). Five homologs from a BLAST search were selected for each enzyme class (**Supplementary Table 1**), synthesized in plasmids, and then amplified as linear expression templates for CFE using *E. coli* cell extracts (**Fig. 1B**). We quantified enzyme expression using ^14^C-leucine incorporation, and the homologs from well-characterized enzyme classes were designated as the “reference pathway” (**Supplementary Fig. 1**). Cell-free reactions containing the reference pathway enzymes at 0.5 µM each, 1 mM NAD, ATP, and CoA, and 100 mM xylose initially produced ~5 mM BT in the context of remaining reagents from gene expression (**Fig. 1B**).

**Figure 1.**
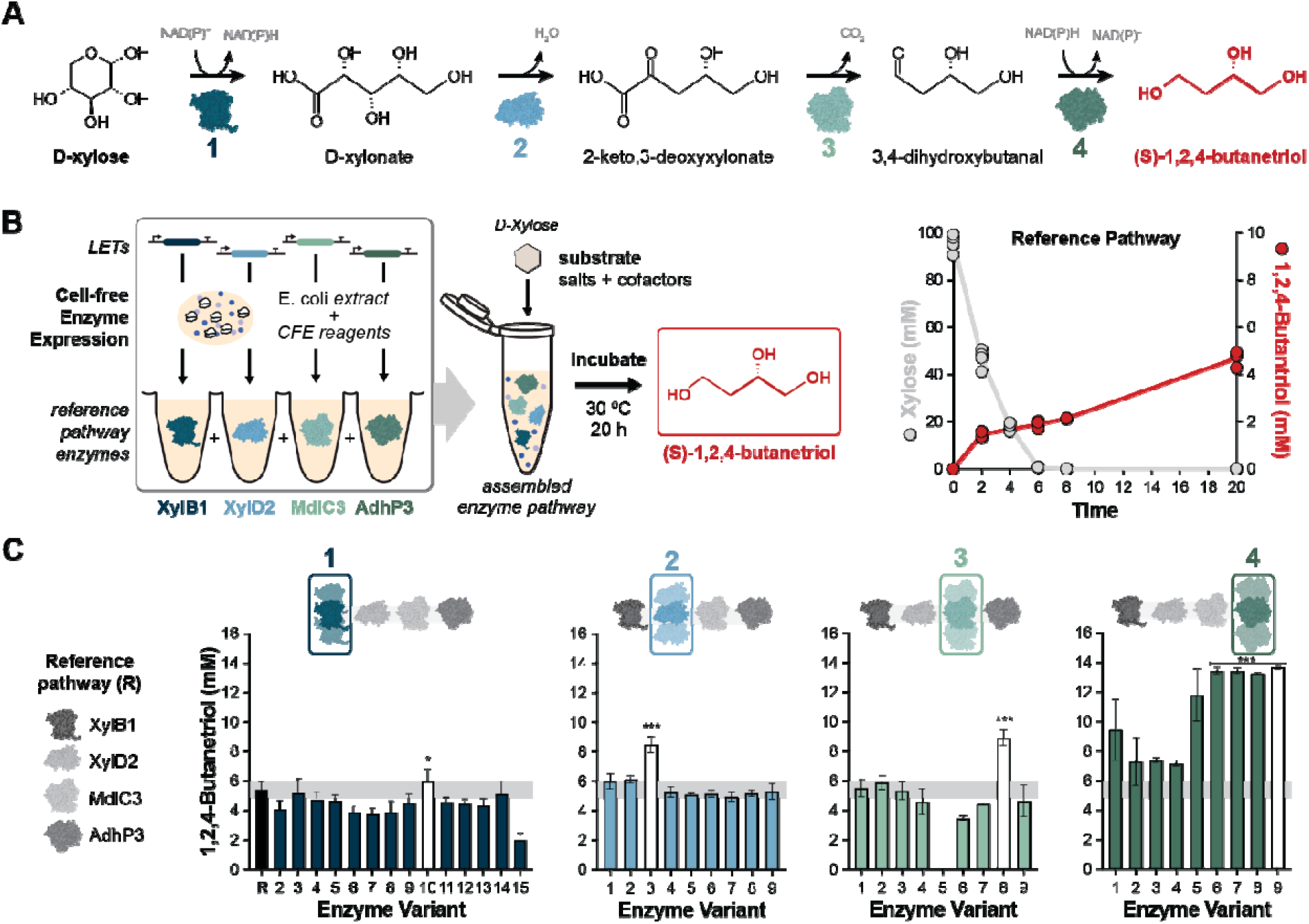
Cell-free gene expression provides a high-throughput framework to rapidly screen and optimize BT titers. The BT synthesis pathway consists of four enzymatic steps from xylose: oxidation, dehydration, decarboxylation, and reduction (A). Enzymes from literature were expressed *in vitro* using an *E. coli* cell extract and combined with xylose, initially producing low BT titers in the context of remaining reagents from gene expression (B). Sequentially replacing each enzyme from the reference pathway with alternative homologs expressed *in vitro* identified better performing enzymes for all four reactions (C). Significance levels indicated by * (p < 0.05) and *** (p < 0.001) are based on two-tailed student’s t-tests of 4 replicates. Data in panel C represent mean ± standard deviation of 4 technical replicates.

With a validated pathway in hand, we sought to increase BT titers by screening the remaining enzyme variants. The reference pathway (R) served as the metabolic backdrop for screening, wherein each reference enzyme was sequentially replaced with alternative homologs at equimolar concentrations to directly assess individual performance contributions (**Fig. 1C**). For the first three steps of the pathway, one homolog performed significantly better than the reference enzyme (Xyd5, XylD4, and MdlC4). The last step of the pathway indicated that all YqhD variants outperform the reference enzyme, more than doubling BT production to 13.7 ± 0.1 mM when AdhP3 was replaced with YqhD5. This follows literature indicating that YqhD has higher activity on 4-carbon substrates than adhP^13,44^. The four best performing enzymes were designated the “improved set” and moved forward for further assessment.

### Increasing BT titers with enriched extracts and an improved decarboxylase

While CFE facilitates rapid enzyme prototyping, previous cell-free metabolic engineering efforts achieved higher conversion efficiencies by using enriched extracts in which heterologous enzymes are expressed *vivo* prior to cell lysis^45^ (**Fig. 2A**). This strategic transition bypasses the metabolic inhibition caused by the reagents required for *in vitro* gene expression, while allowing the loading of significantly higher enzyme concentrations. Enriched extracts containing the reference enzymes and improved enzymes were combined based on bulk protein content (i.e., 5 mg/mL final concentration) to assess conversion of 100 mM xylose, with the reference pathway producing 7.3 ± 0.8 mM BT (**Fig. 2B**). We observed improved BT titers when each of the reference enzymes was replaced with the improved enzyme from the CFE screen, and combining all improved enzymes as enriched extracts produced 100 mM BT (**Fig. 2B**). Then by sequentially omitting each enzyme, we found that *E. coli* background metabolism in the extract retains the capacity to catalyze reactions 1, 2, and 4. We thus identified reaction 3 as the primary rate-limiting bottleneck, prompting a larger exploration of the enzyme space to ensure robust flux through this decarboxylation step^13^.

**Figure 2.**
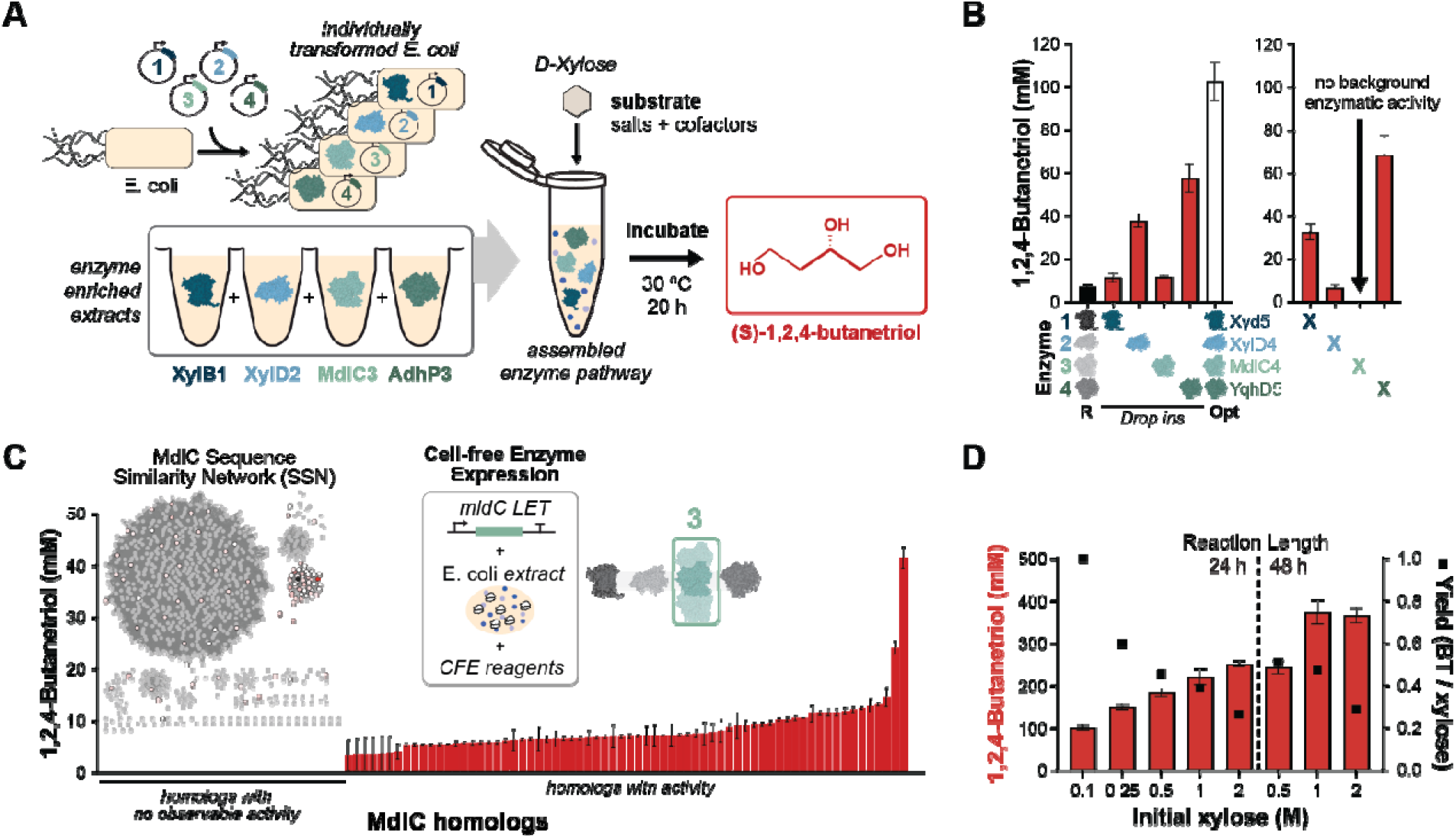
Enriched cell-free extracts maximize BT titers and isolate pathway bottlenecks. Enzymes were expressed *in vivo* before lysis for *in vitro* metabolism (A). The enriched extracts were combined to assess BT production from 100 mM xylose, first replacing each reference enzyme with the improved homolog and then sequentially omitting each enzyme to assess background metabolism (B). Noting no background activity for the third step of the pathway, a larger screening campaign was undertaken using a SSN with ~1000 sequences from a BLAST search of *mdlC4* (C). 96 new homologs were expressed *in vitro* and combined with enriched extracts for steps 1, 2, and 4 to assess BT synthesis (D). The most productive MdlC variant was then used to prepare an enriched extract, which increased BT production from high substrate concentrations. Data represent mean ± standard deviation of 3 technical replicates.

To explore enzyme alternatives for *mdlC4*, the improved enzyme for step 3, we generated a sequence similarity network with ~1,000 sequences from a BLAST search of the enzyme (**Supplementary Table 2**). To compress this large sequence space into a manageable 96-well format, we selected all 46 genes from the cluster containing *mdlC4* and a proportional number of genes from the remaining clusters based on the percentage of total sequences (**Fig. 2C**). Plasmids containing the new *mdlC* variants were synthesized and amplified as linear expression templates for CFE. Adding an equal volume of completed CFE reactions to a mixture of enriched extracts containing Xyd5, XylD4, and YqhD5 enabled screening based on the combination of expression and activity in the MdlC panel (**Fig. 2C**).

Several interesting findings emerged from this screen. Variants producing no detectable BT primarily corresponded to active site mutants with ablated activity, while enzymes in the large cluster with < 65% sequence identity produced up to 12 mM BT (**Supplementary Table 3**). Most importantly for the purpose of maximizing BT synthesis, two variants significantly outperformed the others, corresponding to MdlC variants 35 and 37. Each of these variants has ~97% identity to the source sequence *mdlC3*, representing 13-15 amino acid differences (**Supplementary Table 3**). After ^14^C-leucine incorporation to normalize enzyme loading, both new variants enabled significantly more BT production than the reference enzyme MdlC3, up to 60 mM (**Supplementary Fig. 2**). Formulating MdlC variant 37 as an enriched extract rather than from CFE successfully elevated pathway flux, driving complete conversion of 100 mM xylose to BT and pushing titers past 350 mM BT under high substrate loading (**Fig. 2D**). Yield of BT produced per xylose consumed decreases with higher substrate loading, which is not substantially alleviated when the reaction is extended from 24 to 48 h despite increased BT titers. Substrate inhibition from the large initial concentrations of xylose can be reduced by fed-batch reactions, although increasing the conversion rate requires further reaction optimization (**Supplementary Fig. 3**).

### Developing optimized reaction formulations through design-of-experiments

We next sought to optimize the reaction components to achieve maximum conversion for minimum cost. Rational design of formulations indicated increasing the concentration of NAD while omitting CoA and ATP could improve xylose conversion (**Supplementary Fig 4**). To rigorously account for non-linear interactions, synergistic effects, and blending dynamics between complex reaction components, we carried out a multi-dimensional Design-of-Experiments (DoE) framework to map high-yielding, low-cost formulation spaces. The parameters for optimization included substrate (xylose), buffer (BisTris), cofactors (NAD, CoA, ATP), and the BT pathway enzymes in enriched extracts mixed at an equimolar ratio. Xylose was limited to 500 mM avoid substrate inhibition and enable higher yields (see **Fig. 2D**) in batch mode.

Fifty-five experimental conditions were run in duplicate to estimate component effects and generate predictive models (**Fig. 3A**; **Supplementary Table 4**). This led to the generation of an 18-term regression model and a self-validating ensemble model accounting for linear and non-linear blending effects, both of which aligned well with experimental data and enabled prediction of optimal formulations (**Fig. 3B**; **Supplementary Table 5**). The initial optimization was performed with only BT titer in mind, predicting ~40-50% yields. A secondary optimization accounted for both BT titer and overall cost, predicting ~30-45% conversion with costs ranging from $0.80 to $30 per liter of reagents, excluding enzymes. The predicted formulations from both DoE runs were evaluated at the 10-microliter scale, reaching BT titers from 87 to 182 mM with costs from $0.36 to $300 per gram of product (**Fig. 3C**). The most promising conditions were selected based on minimum cost (formulation #2) at $0.36 / g BT and maximum titer (formulation #24) at 182 mM. Notably, formulation #2 consists solely of xylose and enzyme-enriched cell extract, with reaction co-factors natively present in undialyzed extract^46^. These conditions were subsequently tested with larger volumes to assess the scalability of this BT synthesis platform.

**Figure 3.**
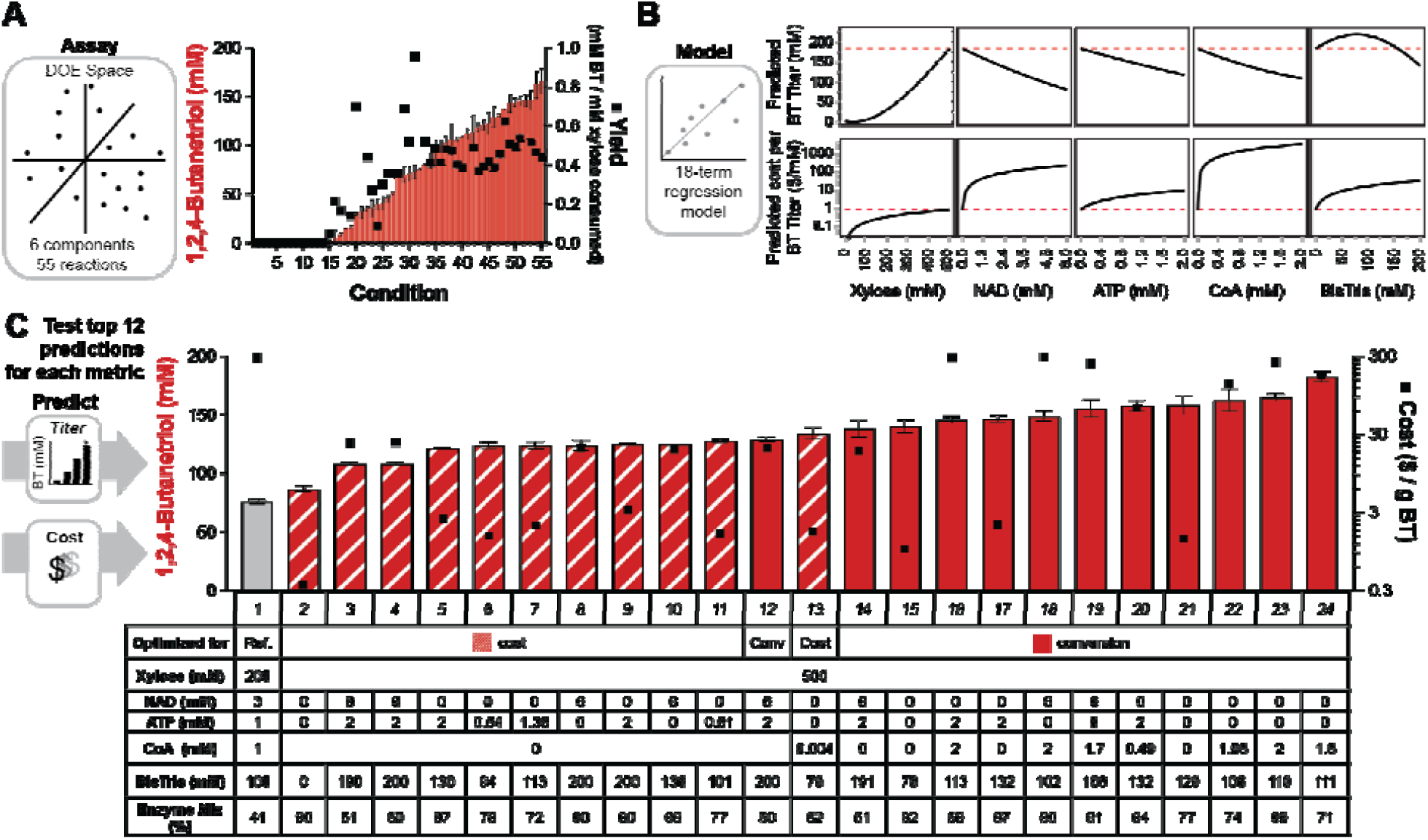
Multi-dimensional design-of-experiments (DoE) accelerates cell-free reaction optimization. A range of test conditions was constructed and run in duplicate, varying 6 components across a wide range of concentration levels (A). The training set informed a regression model to predict formulations that maximize conversion of xylose to BT or minimize reaction costs per gram of product, first modeled with the effects of changing individual components to assess the multidimensional space (B). The most promising predictions were run at the 10 µL scale to compare titer and cost to the reference condition (gray) used in Figure 2, with formulations optimized for cost (striped) or titer (solid red) (C). The minimum cost and maximum conversion conditions (2 and 24, respectively) moved forward for comparison at larger scales. Data represent mean ± standard deviation of 3 technical replicates.

### Scaling from prototyping volumes toward biomanufacturing

After screening enzyme variants and reaction conditions at the 10 µL scale, we sought to establish a bench-scale biomanufacturing setup. Although cell-free reactions dependent on glycolysis typically require large surface-area-to-volume ratios to maintain oxidative phosphorylation,^26,47^ the pathway from xylose to BT appears far less oxygen sensitive. BT titers remained stable in reactions across 3 orders of magnitude, from 10 µL to 1 mL with no significant differences observed with variable headspace volumes (**Supplementary Fig 5**). The process for growing strains expressing the BT pathway enzymes and generating enriched extracts remained constant to the milliliter scale, but pursuing reactions at the 1-L scale required significantly more biomass. The extract preparation protocol was adapted from flask cultivations to bioreactors to facilitate scale-up.

Two formulations from the DoE optimization above were selected for scaling: #2 for minimum cost and #24 for maximum titer. Reactions ran for 48 h in gradually larger vessels, noting that the smallest reactions were set up with separate replicates for each time point while the larger reactions were sampled over the duration of the experiment. The 10 µL and 100 µL reactions used rounded and flat 96-well plates, respectively. The 1 mL and 10 mL reactions used T25 and T150 culture flasks, respectively. The 1 L reactions occurred in “cell factory” chambers with 10 layers to best mimic the airflow and diffusion effects at the smaller scales. Aside from the 100-µL reactions, all conditions achieve at least 10 g/L BT (>94 mM) within 20 h, and most reactions are effectively completed at this point (**Fig. 4**). The 10-mL reactions in T150 flasks performed best, producing 240.8 ± 15.1 mM BT (the highest titer achieved in this study) with the maximum titer formulation and 139.2 ± 3.6 mM BT with the minimum cost formulation after 20 h (**Fig. 4B**). We observed peak productivities of 1.45 g/L/h and 0.91 g/L/h in the max titer and minimum cost formulation, respectively. We chose static batch reactions to isolate the effects of reaction scale as much as possible and to implement low-cost equipment for ease of access in distributed production. Stirring would become essential for homogeneity at commercial volumes, but sparging would likely favor competing reactions in the extract with oxygen dependence. Further optimization and scaling would be best suited to a bioreactor system running in fed-batch mode to maximize BT synthesis and reaction longevity^19^.

**Figure 4.**
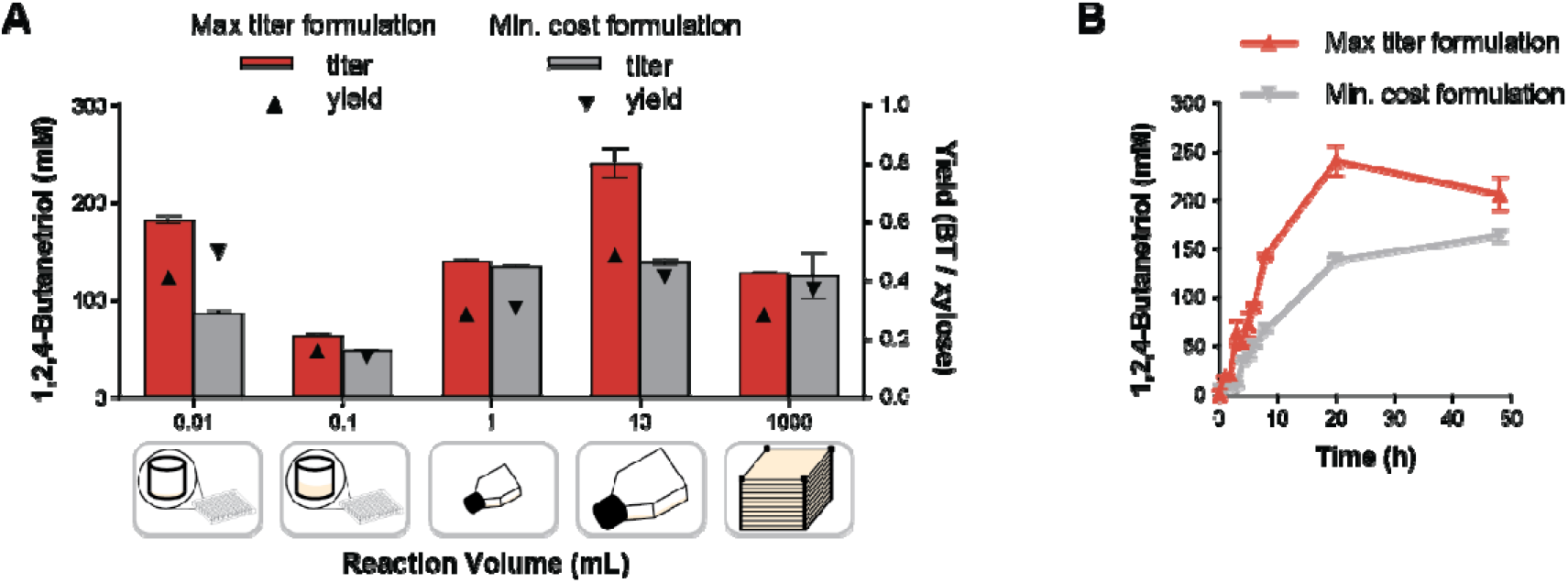
Prototyping to pilot-scale validation of optimized cell-free reactions across five orders of magnitude. The best-performing formulations for minimized cost and maximized yield were tested across 5 orders of magnitude from 10 µL to 1 L, showing average titer and yield at 20 h (A). The 10 mL setup best reflected the 0.01 mL prototyping scale with higher titers shown in a representative time course (B). Data represent mean ± standard deviation of 3 technical replicates.

## Discussion

In this work, we establish an end-to-end cell-free paradigm for discovery, development and manufacturing of platform chemicals within a single, programmable framework. By bypassing living cellular constraints, the entirely cell-free architecture holds promise to compress the time to product and avoid iterative efforts in cells. As a model pathway, we explored the production of the platform chemical BT.

To establish this platform, we systematically studied nearly 150 enzyme variants for BT synthesis from xylose in a cell-free system, screening with CFE before moving toward biomanufacturing with enriched extracts. After optimizing reaction conditions with the best homologs for each step of the pathway using DoE, we scaled reactions five orders of magnitude, from 10 microliters to 1 liter, producing >10 g/L BT in all volumetric scales. This establishes the purification-free, enriched extract setup as a viable biosynthesis platform for chemical manufacturing, with reagent costs in the minimized formulation totaling just $3.00 per liter (the cost of the raw substrate xylose). We primarily explored natural homologs of BT enzymes, including distant alignments with <60% sequence identity that still showed some activity in the third reaction. Deeper exploration of the sequence space through targeted mutagenesis and machine learning-guided engineering has the potential to further increase conversion rates and efficiencies, as demonstrated in campaigns for amide synthetases^39^ and glycoside hydrolyases^48^. The convergence of enzyme stability against temperatures and solvent exposure could greatly enhance BT biomanufacturing using the enzymes validated here as scaffolds for engineering, enabling faster reactions rates with elevated temperatures and aiding *in situ* extraction of BT during synthesis^19,49,50^.

The cell-free biosynthesis strategy delivers a streamlined, design-driven architecture for the scalable, cost-effective production of BT (maximum titer of 39.6 g/L in this study; **Fig. 2**) (**Supplementary Fig. 6**). While purified multi-enzyme cascades exhibit high conversion capacity (up to 170 g/L), the prohibitive cost of affinity chromatography and exogenous cofactor supplementation renders purified systems economically unviable for commodity chemicals^14^. Conversely, while cellular fermentations have demonstrated titers of BT from xylose exceeding 40 g/L^15,51^, living hosts rely on complex media that can confound yield calculations^52^ and rely on large amounts of base for pH maintenance^4^, which is avoidable in a cell-free system by simply buffering the reaction^53^. We anticipate the future optimization efforts could take cell-extract based bioconversions into commercially competitive regimes. First, implementing fed-batch or semi-continuous reactions for gradual substrate additions could push the optimized cell-free setup toward higher titers, as observed in CFE^54^ and as demonstrated with a version of our BT synthesis reactions (**Supplementary Fig. 3**). Second, while BT and pathway intermediates are stable in the cell extract (**Supplementary Fig. 7**), removal of negative effectors through genomic deletions, protease tagging^55^, affinity purification^56^, or heat inactivation^19^ could be beneficial. Third, as cell-extract preparation scales toward high-density continuous fermentation on defined media, catalyst generation costs will drop dramatically from current batch-scale estimates (~ $1,000/L). Additionally, the 4 enriched extracts could be reduced to a single extract through co-culture and processing with an all-in-one approach that avoids co-expression of the full pathway in a single strain^57,58^. This could result in similar BT synthesis to the 4 mixed extracts with less material and effort in upstream processing (**Supplementary Fig. 8**). Efficient separation from broth will also be critical for large-scale purification, which can be achieved with recyclable boronic acid-functionalized polymers^50^ or potentially via simple phase separation at high enough titers^19^.

Ultimately, this work bridges the long-standing divide between accelerated cell-free discovery and scale-up biomanufacturing. By combining rapid DBTL prototyping with scalable, high-yield chemical synthesis, this platform establishes a fundamental blueprint for sustainable biomanufacturing, especially when in situations where a higher tolerance to toxic compounds or pH variations introduced by pretreated lignocellulosic feedstocks compared to living cells^22^ may be needed. Taken together we anticipate that our work will contribute to the diversification of the biomanufacturing sector as we work toward more efficient and sustainable chemical production strategies at industrial scales.

## Materials & Methods

### Chemicals and DNA

All chemical reagents were supplied by Sigma-Aldrich unless otherwise noted. DNA oligonucleotides were supplied by IDT, and plasmids were synthesized by Twist Bioscience in the pJL1 backbone (Addgene #102634) for T7-driven expression.

### Cell Extract Preparation (prototyping scale)

General protocols and methods were as described previously^59,60^.

#### For cell-free gene expression

*E. coli* strain BL21 Star^TM^ (DE3) was inoculated into LB media with a single colony picked from a freshly streaked plate. After overnight incubation at 37 °C, 1 L of 2xYTP media (16 g/L tryptone, 10 g/L yeast extract, 5 g/L NaCl, 7 g/L of K_2_HPO_4_, and 3 g/L KH_2_PO_4_ adjusted to a final pH to 7.2) in a 2.5 L Tunair flask was seeded with an initial OD_600_ of ~0.075. The cultures were grown at 37°C with shaking at 250 rpm until reaching OD_600_ values between 0.5-0.6 when T7 RNA polymerase expression was induced by adding 200 µL of 1 M IPTG. Cells were harvested in mid-exponential phase upon reaching an OD_600_ of 3.0-3.5.

#### For enzyme enrichment

Chemically competent BL21 Star^TM^ (DE3) cells (New England Biolabs) were transformed with a pJL1 expression plasmid and grown in LB media with 50 µg/ml kanamycin. 1 L cultures in 2xYTP were inoculated at an initial OD_600_ of ~0.075, induced with 200 µL of 1 M IPTG at OD_600_ 0.6-0.8, and harvested after 4-5 hours of enzyme expression (reaching OD_600_ 5-6).

Cells were harvested by centrifuging at 8000 x *g* for 5 min at 4°C, and the pellets were immediately transferred to two 50 mL conical tubes on ice. The cell pellets were washed with 25 mL of cold S30 buffer (50 mM Tris base, 60 mM potassium glutamate, and 14 mM magnesium glutamate, adjusted to pH 7.7) and resuspended through cycles of 15 seconds of vortexing and 15 seconds on ice. The cell suspensions were centrifuged at 10,000 x *g* for 2 min at 4°C, and the supernatant was carefully discarded. The wash procedure was repeated for a total of 3 times before the cell pellets were flash frozen in liquid nitrogen and stored at −80 °C. Prior to lysis, the cell pellets were allowed to thaw on ice for 1 hour and resuspended with 1 mL of S30 buffer per gram of cell pellet. The cells were lysed in a single pass using an Avestin EmulsiFlex-B15 homogenizer at a pressure of 20,000-25,000 psi. The lysed cells were immediately transferred to clean 1.5 mL microcentrifuge tubes on ice. The tubes of lysate were centrifuged at 12,000 x *g* for 10 minutes at 4°C. The supernatant (cell extract) was aliquoted, flash frozen in liquid nitrogen, and stored at −80 °C until use. A Bradford assay was performed to determine the bulk protein concentration of the extract using a bovine serum albumin standard curve.

### Enzyme Variant Generation and Selection

Initial homologs were identified by BLAST searches of the most commonly used enzymes in the pathway from xylose to butanetriol. Homologs indicated with the number 1 were used as the source sequence, and the remaining homologs were at least 90% similar to the source enzymes (**Supplementary Table 1**). NAD(H) and NADP(H)-dependent homologs were identified for screening steps 1 and 4 of the pathway. The sequence similarity network (SSN) for further *mdlC* screening was generated with the EFI Enzyme Similarity Tool^61^ after collecting ~1,000 homologs from a Uniprot BLAST search of *mdlC4* and plotted in Cytoscape with an alignment score threshold of 275 (**Supplementary Table 2**). The SSN was pared down to 96 homologs to facilitate screening via cell-free expression and HPLC, using all 46 variants from the cluster containing *mdlC4* (>85% sequence similarity) and the proportional number of sequences from the remaining clusters (**Supplementary Table 3**).

### Cell-Free Gene Expression (CFE)

Enzymes were expressed using the PANOx-SP formulation for cell-free gene expression as previously described^60^ (8 mM magnesium glutamate, 10 mM ammonium glutamate, 130 mM potassium glutamate; 1.2 mM ATP; 0.85 mM GTP, UTP, and CTP; 0.034 mg/ml folinic acid, 33.3 mM phosphoenolpyruvate, 2 mM of the canonical amino acids, 0.33 mM NAD, 4 mM oxalic acid, 1 mM putrescine, 1.5 mM spermidine, and 57 mM HEPES buffer with 1.33 µL linear expression template (LET) and 2.67 µL *E. coli* extract (~15 mg/ml final concentration) and nuclease free water up to 10 µL final volume). Reactions were supplemented with 200 µM thiamine pyrophosphate for expression of 2-keto acid decarboxylase variants to ensure proper folding and activity. Enzyme expression was quantified based on incorporation of 10 µM ^14^C-leucine with analysis on a PerkinElmer MicroBeta^2^. All reactions were incubated for 20 h at 30°C. LETs were amplified from a pJL1 plasmid backbone encoding each pathway enzyme using Q5 DNA polymerase (NEB) with standardized primers^60^.

### Cell-Free 1,2,4-Butanetriol Synthesis

10 µL reactions were prepared in 1.5 mL microcentrifuge tubes to screen different combinations of enzymes from each step of the BT production pathway. Each butanetriol synthesis reaction comprised the following reagents unless otherwise specified: 8 mM magnesium glutamate, 10 mM ammonium glutamate, 134 mM potassium glutamate, 100 mM BisTris Buffer, 3 mM NAD, 1 mM ATP, 1 mM Coenzyme A (CoA), 100 mM to 2 M D-xylose, 1 µM of each expressed enzyme from CFPS or 5 mg/mL of each enzyme-enriched extract from enriched extract. Reactions were incubated at 30°C for 20 h. After screening enzyme variants using this formulation, the composition was optimized as described through DOE.

### Metabolite Analysis

Cell-free reactions were quenched with an equivalent volume of 10% w/v trichloroacetic acid. The samples were centrifuged for 10 min at 20,000 x *g* and the resulting supernatant in each 1.5 mL microcentrifuge tube was transferred to a vial for HPLC analysis. The samples were run on a Phenomenex Rezex ROA-Organic Acid H+ column (catalog number: 00H-0138-K0) with a mobile phase of 5 mM sulfuric acid and a flow rate of 0.6 mL/min.

### Design-of-Experiments (DoE)

JMP^®^ Pro 18 software was used to assemble a DoE design based on the Scheffé class of mixture designs. It was a 55-sample hybrid design that used 25 Space Filling points and 30 I-Optimal points (**Supplementary Table 4**). The hybrid (I-Optimal/Space Filling) design along with the special cubic mixture model was designed to detect linear-blending effects as well as any 2-way or 3-way synergies (non-linear blending effects) that may exist among combinations of the formulation components xylose, NAD, ATP, CoA, BisTris, and enzyme mix (enriched extracts at an equimolar ratio). The starting model had 41 terms and included linear blending, two-way, and three-way non-linear blending terms. The response data from the 55 DOE samples was used to fit two separate prediction models for BT titer: (1) an 18-term reduced regression model created with Design-Expert 25 software using forward step-regression with an AiC stopping criteria, (2) a Generalized Regression based self-validated ensemble model (SVEM) created with JMP® Pro 18 using Lasso (penalized) regression with an ensemble of 200 intermediate models. The SVEM model and the 18-term reduced regression model were each used to predict optimal formulations to maximize BT titer. Finally, the BT titer prediction models were used in conjunction with a deterministic formulation cost function to perform a multi-objective optimization aimed at simultaneously maximizing BT titer while minimizing formulation cost. Additional details of the design and validation are provided in **Supplementary Note 1**.

### Large Scale Extract Preparation

In general, cell-free lysates were produced following the protocol in the cell extract preparation for enzyme enrichment section. When necessary, the protocol was modified as follows to accommodate the scale of the preparation:

*E. coli* strain BL21 Star^TM^ (DE3) containing plasmids expressing each enzyme were plated on fresh 2xYTP agar plates and incubated overnight at 37°C. Individual seed cultures of 1 L 2xYTP with 50 µg/mL Kanamycin were started 16 hours before the final culture was inoculated. A 50L (New Brunswick, BioFlo 610) bioreactor containing Overnight TB express medium (Novagen) was inoculated with seed cultures at an initial OD_600_ of ~0.075. The fermenter settings were adjusted to 300 rpm, 50 SLPM air flow and the dissolved oxygen (DO) meter was calibrated to 100%. Cells were harvested after 12 hours of enzyme expression (OD_600_ ~10). Each enzyme-enriched lysate was grown and processed separately.

Cells were harvested by centrifuging using a pre-chilled (4°C) Powerfuge Pilot 1.1 L bowl system (CARR Biosystems) at 10,000 x *g* at 4°C, and the pellets were scraped from the bowl and immediately transferred to plastic freezer bags, weighed, flattened, and flash frozen. The

cell pellets were removed for processing at least 24 hours after they were frozen, resuspended in cold S30 buffer (50 mM Tris base, 60 mM potassium glutamate, and 14 mM magnesium glutamate, adjusted to pH 7.7) at a 1 mL per gram of pellet ratio. The cell suspensions were lysed in a single pass using an M110P Microfluidizer Processor at a pressure of 20,000-25,000 psi. The lysed cells were immediately transferred to clean 250 mL centrifuge bottles on ice. Centrifugation, flash freezing, and storage of the large-scale extracts from this point match the protocol above for prototyping scale preparation.

### Large Cell-Free Butanetriol Synthesis

Progressively larger reaction vessels were used to run larger reactions to approximate similar ratios of surface area to volume. 10 µL reactions were run in round-bottom 96-well plates, 100 µL reactions in flat-bottom 96-well plates, 1 mL reactions in T25 culture flasks, 10 mL reactions in T150 flasks, and 1 L reactions in 10-layer Cell Factories (Nunc). All reactions were composed of the same reagents across scales and concentrations of expressed enzymes were held at 5 mg/mL based on Bradford measurements.

## ACKNOWLEDGEMENTS

We are grateful to Corinne Scown and Tyler Huntington for insights into the Bio-C2G platform for technoeconomic analysis. This work was supported by Army Contracting Command (W52P1J-21-9-3023) and the Army Research Office (W911NF-22-2-0246).

## Author Contributions

Conceptualization: B.J.R., M.C.J., A.S.K., M.W.L.

Investigation: B.J.R., K.A.R., I.D.R., J.T.L., M.S.L., D.C.G., K.Z., J.D., E.N.W., J.R.B., F.J.K.

Writing – Original draft: B.J.R.

Writing – Review and editing: K.A.R., I.D.R., M.W.L., A.S.K., M.C.J.

Visualization – B.J.R., A.S.K., J.T.L., K.Z.

Project administration: M.C.J., A.S.K., M.W.L.

Funding acquisition: M.C.J., M.W.L.

