## Supplemental Figures for "Cell-free pathway prototyping enables cost-effective biomanufacturing of 1,2,4-butanetriol at the 1-L scale"

1. Quantification of cell-free gene expression
2. Fed-batch reaction data
3. Titration of enzymes from MdlC screen
4. Mixed extracts vs all-in-one
5. Cofactor titration
6. Test of vessels and scales
7. TEA snapshot
8. Metabolite stability

**Supplemental Tables (see Excel file)**

1. Sequences and accessions of initial enzymes
2. Sequences in MdlC SSN
3. Sequences of enzymes screened from SSN
4. DOE training data table
5. DOE results

**Supplemental Figures**

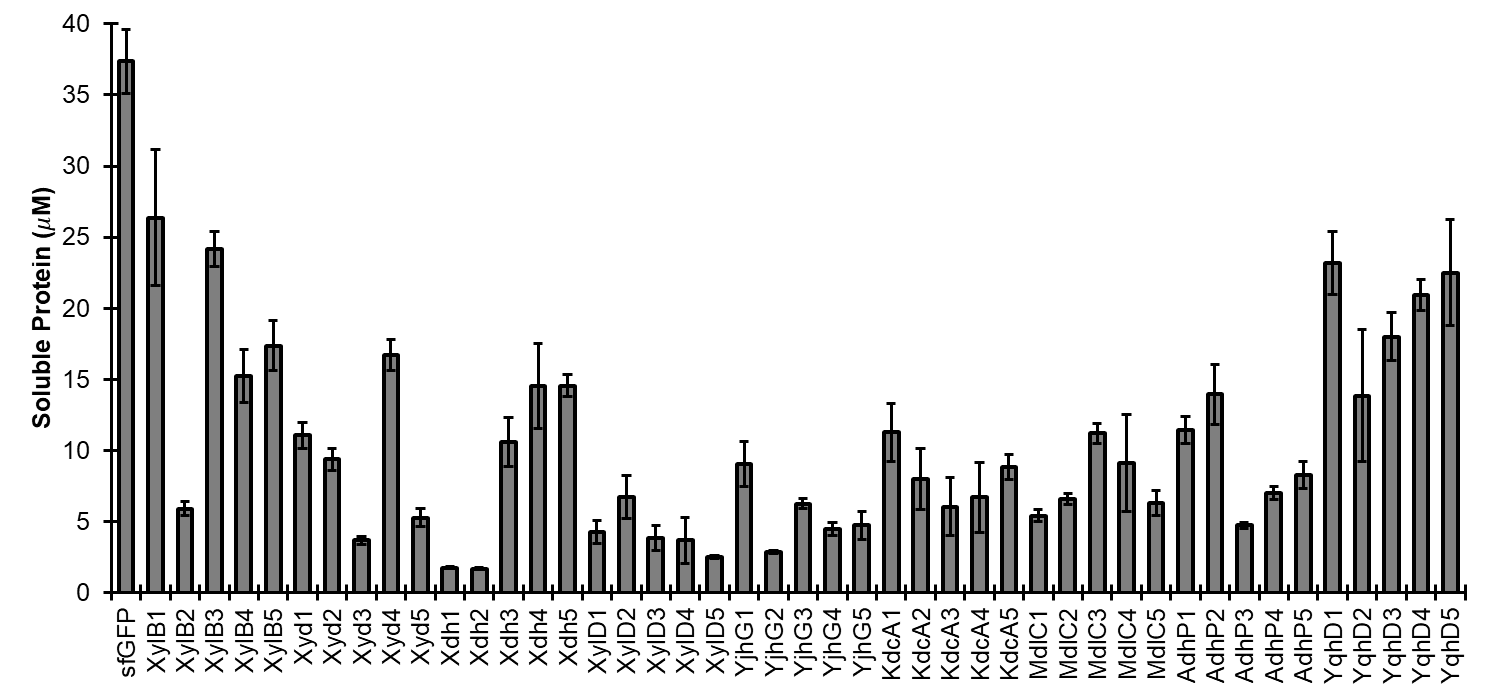

**Supplementary Figure 1. Quantification of cell-free expression by ^14^C-leucine incorporation.** Linear expression templates were transcribed and translated in cell-free reactions containing isotopically labeled leucine, and the soluble fraction was quantified to compare equimolar amounts of homologs for each step of the pathway. Super-folder green fluorescent protein (sfGFP) serves as a highly expressed positive control.

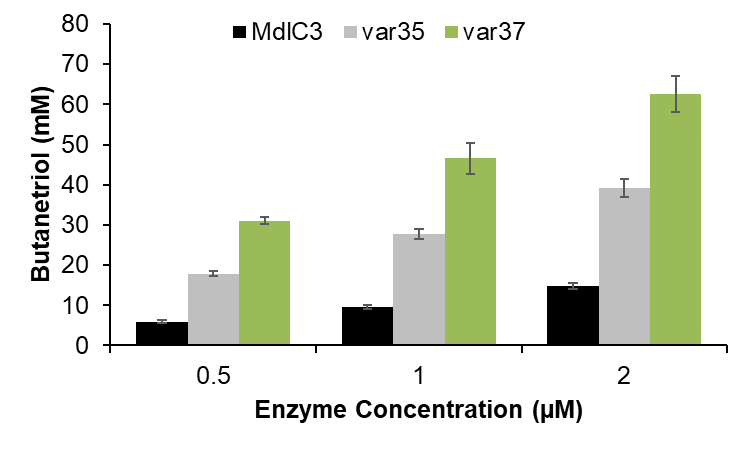

**Supplementary Figure 2. Titration of enzymes from Sequence Similarity Network (SSN).** At a range of concentrations, the top performing enzymes from the SSN screen continue to produce higher BT titers than the initial MdlC variant from the reference pathway in **Figure 1** (see main manuscript).

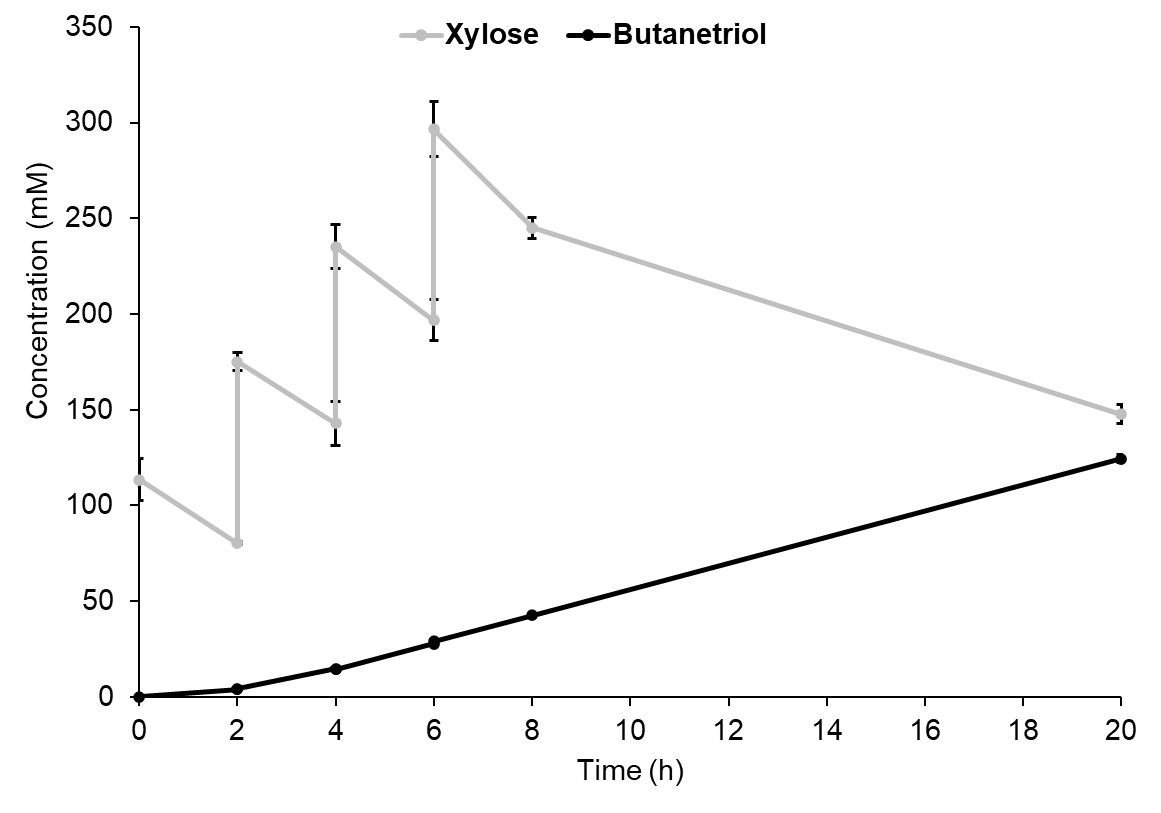

**Supplementary Figure 3. Fed-batch reaction data.** Supplementing 100 mM xylose in 2-hour increments slows butanetriol synthesis and results in relatively low conversion. Extended reaction times or more continuous substrate additions could alleviate substrate inhibition.

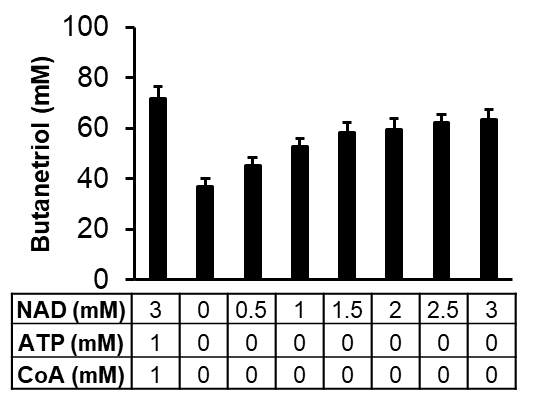

**Supplementary Figure 4. Cofactor optimization.** ATP and CoA are unnecessary for these reactions, and no additional NAD still enables ~50% of the titer compared to 3 mM NAD with 200 mM initial xylose.

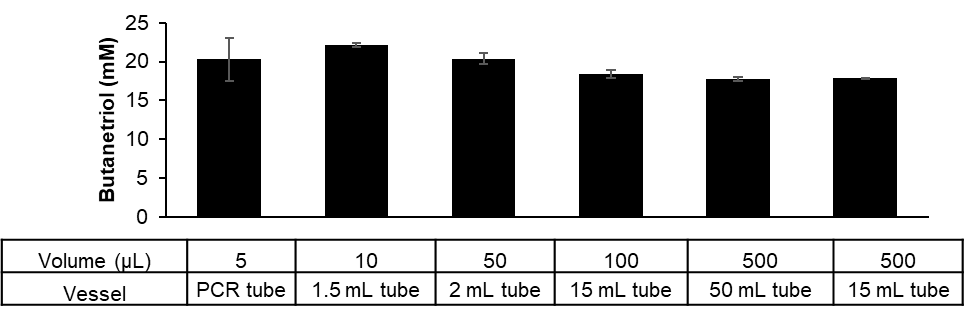

**Supplementary Figure 5. Initial scaling assessment.** A range of reaction volumes from 5 to 500 µL shows small impacts from the ratios of surface area to volume or liquid to headspace in the tube.

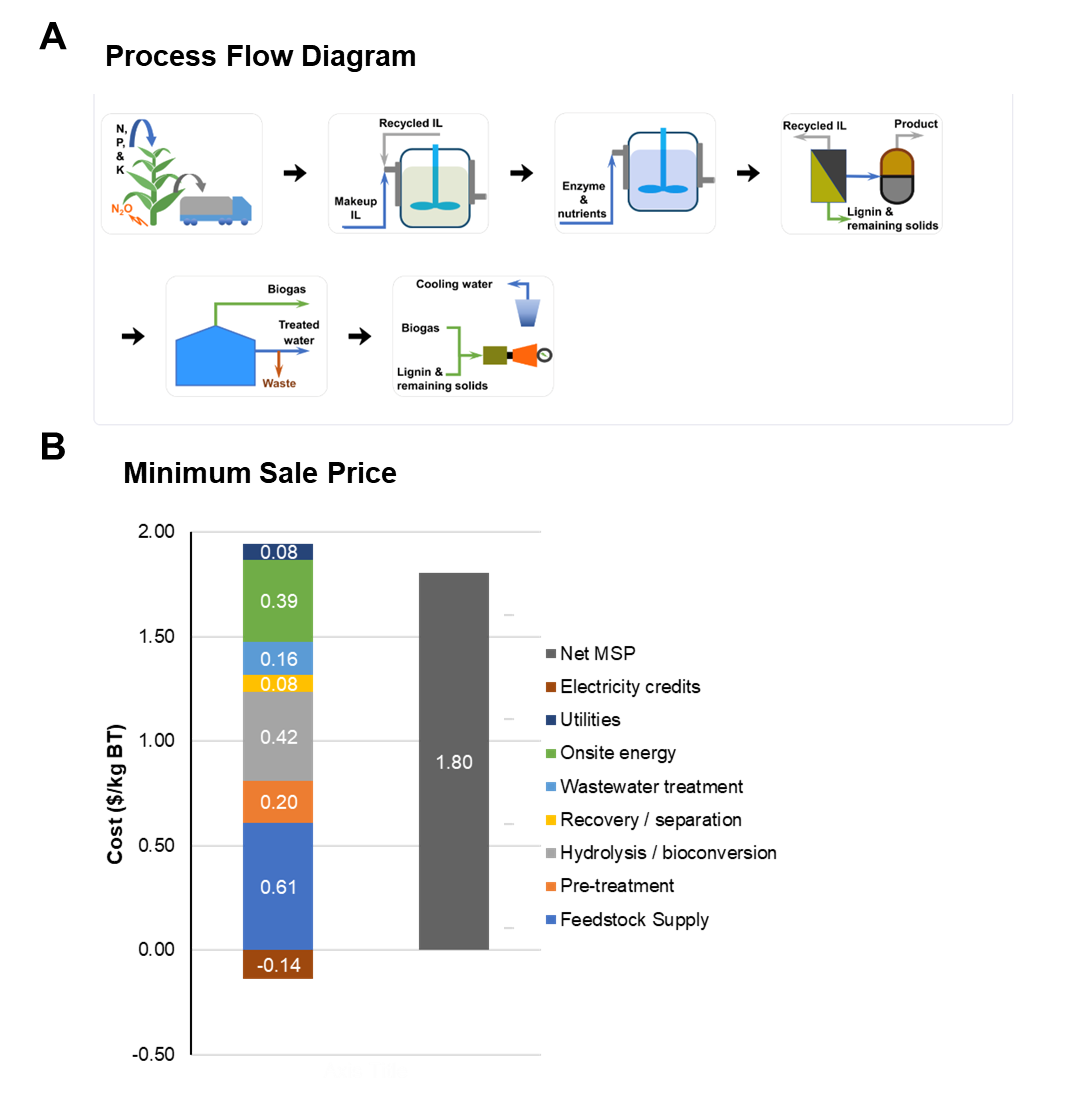

**Supplementary Figure 6. Technoeconomic Analysis.** **(A)**The BioC2G program was used to model a large-scale operation converting xylose from lignocellulosic biomass into butanetriol using enzymes. **(B)** The minimal sale price was predicted to be $1.80 per kilogram of butanetriol to break even with annual capital and operating expenses. Default parameters were used, including the need to process 2000 metric tons of feedstock per day.

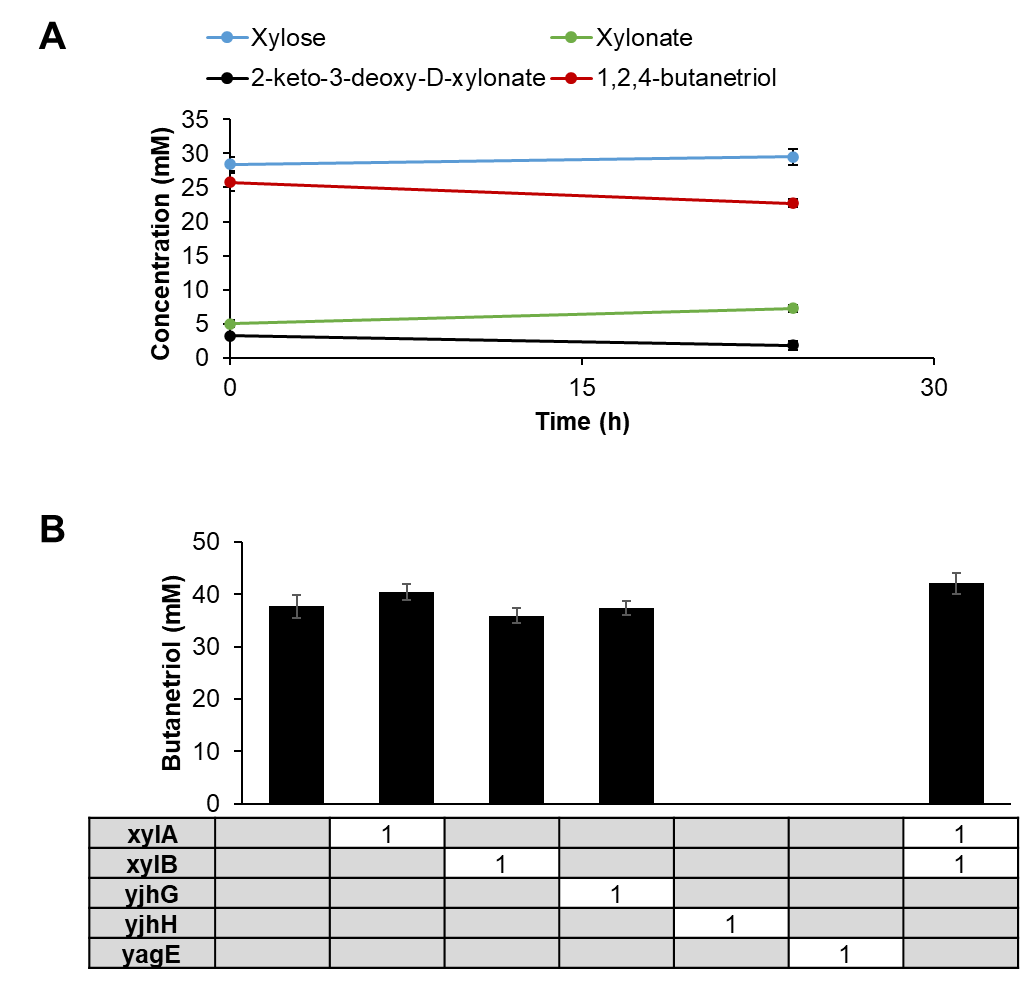

**Supplementary Figure 7. Metabolite stability.** **(A)** The commercially available pathway intermediates remain stable in *E. coli* extract over a 20-hour incubation. **(B)** Supplementing 1 µM of endogenous enzymes that could influence the conversion of xylose to butanetriol indicates potential competition from yjhH and yage. However, based on the stability of intermediates in extract, it appears that these enzymes are not significantly expressed under the conditions at which *E. coli* is harvested prior to extract preparation.

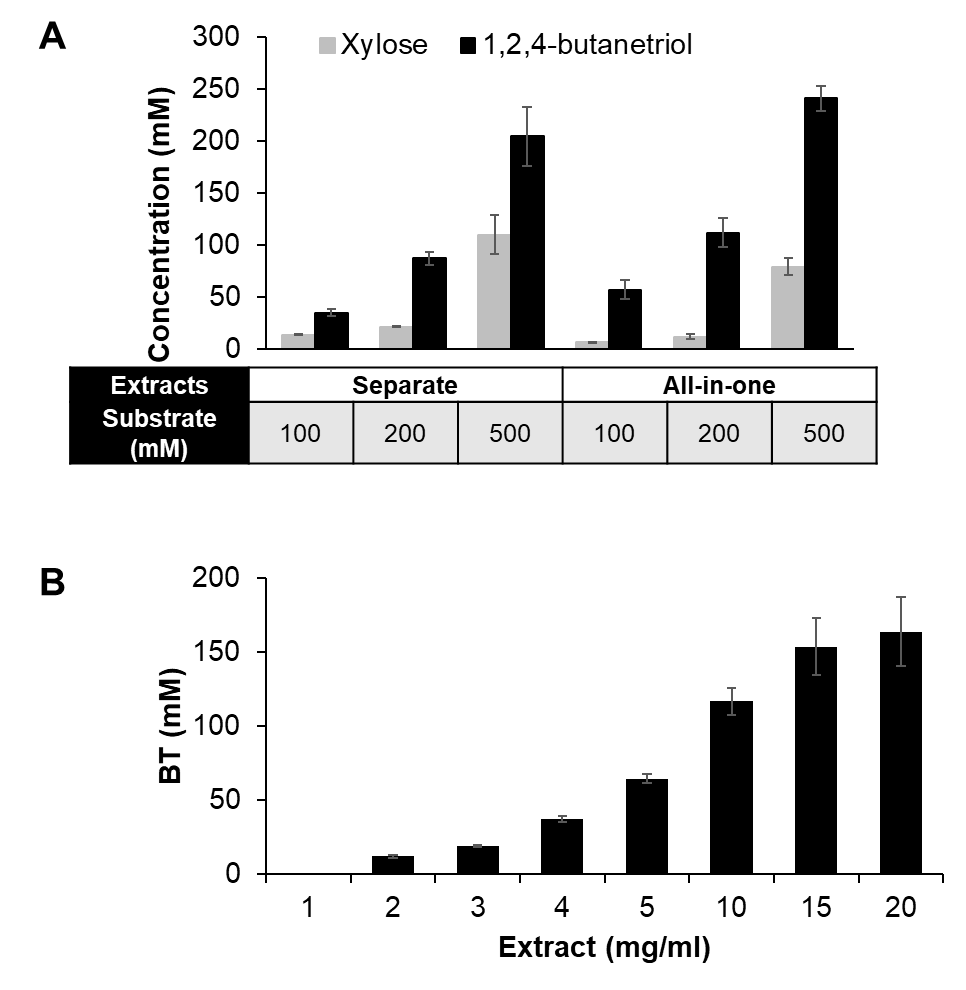

**Supplementary Figure 8.** **Mixed extracts can simplify processing.** (A) Reactions containing equal parts of enriched extracts from 4 separate cultures perform similarly to extracts prepared from a co-culture of 4 strains expressing the BT pathway enzymes. (B) Preparing and titrating a single extract could simplify scale-up efforts.
